# Beyond bold versus shy: Zebrafish exploratory behavior falls into several behavioral clusters and is influenced by strain and sex

**DOI:** 10.1101/2022.04.11.487928

**Authors:** Neha Rajput, Kush Parikh, Justin W. Kenney

## Abstract

Consistent individual differences in exploratory behavior have been found across a range of taxa, including zebrafish, and are thought to contribute to evolutionary fitness. Animals that explore more of a novel environment, and visit areas of high predation risk, are considered bold, whereas animals with the opposite pattern of behavior are considered shy. Here, we examined whether this bimodal characterization of bold versus shy adequately captures the breadth of exploratory behavior exhibited by zebrafish or if, instead, behavior falls into multiple distinct behavioral subtypes. To identify the presence of behavioral subtypes, we applied unsupervised machine learning to behaviors extracted from three-dimensional swim traces from over 400 adult zebrafish across four strains (AB, TL, TU, and WIK) and both sexes. We found that behavior stratified into four distinct clusters. These included previously described bold and shy behavior as well as two new behavioral types: wall-huggers and active explorers. Consistent with prior work that found individual differences to be stable across time and influenced by biological factors like genetics and sex, the behavioral subtypes we identified were stable for up to 10 weeks and were influenced by the strain and sex of the animals. Taken together, our work suggests that individual differences in zebrafish exploratory behavior goes beyond bold versus shy and exhibits greater complexity than is typically assumed.

## Introduction

Interest in the biology of behavioral differences can be traced back to at least Roman antiquity where Galen applied the humoral theory to explain variation in human temperament (as translated in Grant, 2000). Today, we have a greater understanding of human personalities, defined as behavioral tendencies that are consistent across time and context, but its biological basis remains elusive. One avenue for progress is the modeling of human personality through the study of individual difference in animal behavior. Often dismissed as noise around an average, a growing body of work has found that variations in animal behavior are often consistent across time and context (Dall et al., 2012; Sih et al., 2004). Such consistent differences have been described for behaviors important for evolutionary fitness, and in a wide range of taxa, suggesting that they may provide grist for adaptation to an ever-changing environment.

One of the most widely studied axes of behavioral variation in animals is the bold-shy axis. Bold animals tend to explore or investigate novel environments or objects more readily than shy animals that tend to flee or retreat in response to novelty (Réale et al., 2007; Toms et al., 2010; Wilson et al., 1994). From a fitness perspective, boldness may be adaptive when food resources are scarce and predation risk is low, whereas shyness may be more effective when the opposite conditions prevail (Réale et al., 2007). Variation along this axis has been described in animals ranging from bears (Myers and Young, 2018), lizards (López et al., 2005), birds (Carere et al., 2005), and fish (Toms et al., 2010). Studies examining the bold-shy axis typically begin with the assumption that animals fall into one of these two categories. However, whether this bimodal distribution of bold versus shy fully captures animal behavior has not been tested directly. It is likely that there is more complexity to animal behavioral types which may have been overlooked due to use of small sample sizes or assessment of only one or two specific behaviors (Forkosh et al., 2019).

Zebrafish have proven to be an excellent model organism to understand behavior and its biological basis. With 70% of fish genes having an obvious human ortholog (Howe et al., 2013), and a central nervous system that has the same general organization and utilizes many of the same neurotransmitters as mammals (Kenney et al., 2021; Panula et al., 2010; Wulliman et al., 1996), findings using zebrafish are widely applicable. Our understanding of zebrafish behavior has expanded rapidly over the past decade (Gerlai, 2020; Kalueff et al., 2013; Kenney, 2020), including several studies that have examined the bold-shy axis. Boldness is often probed by exposing fish to a novel tank and examining locomotion, exploratory activity and avoidance behaviors, like geotaxis (i.e. bottom dwelling) or thigmotaxis (i.e. proximity to tank walls) (Mustafa et al., 2019; Oswald et al., 2012; Thörnqvist et al., 2019; Toms et al., 2010). Animals that are more active or spend more time in parts of the tank that would increase risk of predation (i.e. the top and/or center of the tank) are considered bolder. In zebrafish, these behaviors have been found to be consistent over time (Baker et al., 2018; Tran and Gerlai, 2013) and can predict other behaviors like social dominance (Dahlbom et al., 2011), aggression (Martins and Bhat, 2014), and stress reactivity (Oswald et al., 2012), all hallmarks of personality. However, fully utilizing zebrafish exploratory behavior to understand the biological basis of individual differences requires that we first determine if the bold versus shy distinction adequately captures the breadth of behavioral variability exhibited during exploration.

To determine the presence of multiple behavioral clusters during exploration of a novel tank we captured three-dimensional swim behavior from over four hundred fish. Because zebrafish behavior is known to be influenced by strain and sex (Volgin et al., 2019), we used animals from four inbred strains (AB, TU, WIK, and TL) and both sexes to ensure that we captured the full range of behavioral variability. Using an unsupervised machine learning approach to identify behavioral clusters we found that exploratory behavior stratified into four clusters. These clusters included traditional descriptions of bold and shy, as well as two additional behavioral types we call wall-huggers and active explorers. Consistent with these behavioral subtypes being akin to personality types, we found that individual cluster membership remained consistent across days and weeks, and that the proportion of fish in each cluster was influenced by strain and sex.

## Results

### Three-dimensional behavioral tracking

To capture three-dimensional zebrafish swim behavior during exploration of a novel tank, we used Intel RealSense^TM^ cameras mounted above five-sided tanks with frosted walls (Figure 1A). These cameras simultaneously captured color and depth streams, resulting in three-dimensional videos (Figure 1B). Fish posture at each frame was tracked in the color stream using DeepLabCut (Figure 1C; Mathis et al., 2018). These points were overlayed onto the depth stream to create three-dimensional swim traces (Figure 1D) from which we extracted positional information (distance from bottom and center), distance travelled, and percent tank explored (Figure 1E).

**Figure 1.**
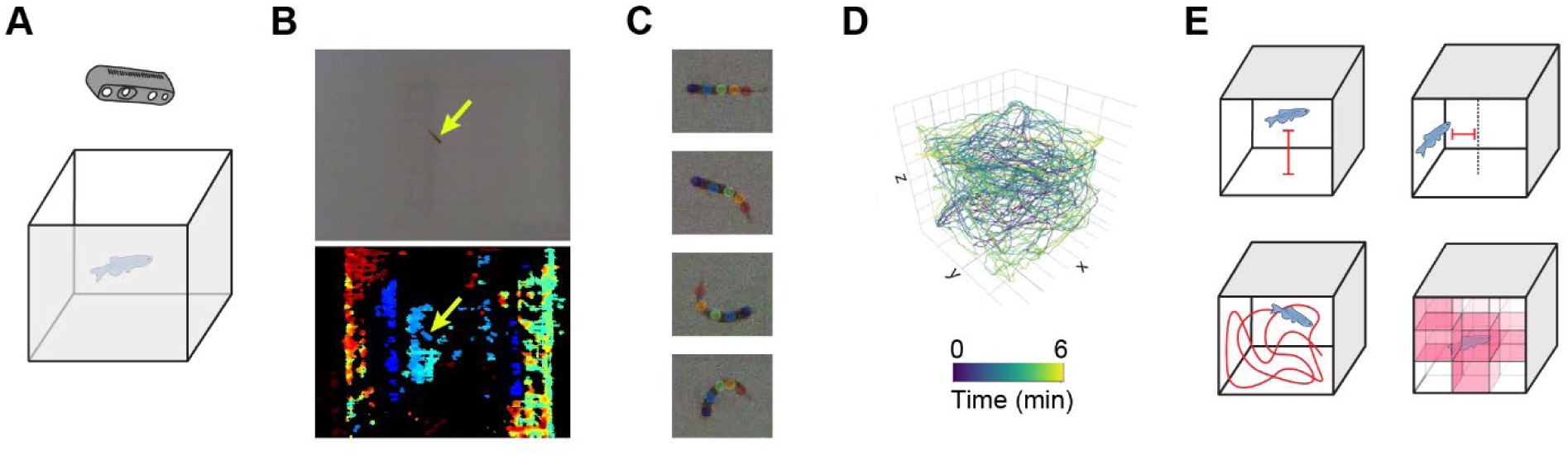
Overview of three-dimensional behavioral tracking. **A**) Individual fish were placed into a novel tank while video was recorded from above using Intel RealSense^TM^ cameras. **B**) Videos included both a color (top) and a depth (bottom) stream where fish can be seen (yellow arrows). **C**) Animals were tracked in the color videos using DeepLabCut to identify five points along the length of the fish. **D**) Tracking was overlaid with the depth stream to generate a three-dimensional trace for each animal. **E**) Four exploratory parameters were extracted from each trace: bottom distance (top left), center distance (top right), distance travelled (bottom left), and percent of the tank explored (bottom right).

### Influence of sex and strain on exploratory behaviors

We first determined if sex and genetics influences individual zebrafish exploratory behaviors by assessing swim traces of fish from four strains (AB, TU, TL, and WIK), and both sexes, on two consecutive days. We extracted four exploratory behaviors from each swim trace: distance from bottom, distance from center, total distance travelled, and percentage of tank explored (Figure 1E). We found that the distribution of several of the parameters deviated from normality (Figure S1), so we performed non-parametric 4 × 2 (strain × sex) permutation ANOVAs. For distance from bottom (Figure 2A), we found a main effect of strain (P = 0.0001), but no effect of sex (P = 0.40) nor an interaction (P = 0.44). FDR (false discovery rate) corrected permutation t-tests found that TL fish swam closer to the bottom of the tank than all other strains (P’s ≤ 0.0008). For distance from center (Figure 2B) there was a main effect of sex (P = 0.042) and a trend towards an interaction (P = 0.060) where female fish of every strain, except AB, spent more time closer to the center of the tank. We also found a main effect of strain (P = 0.0001) with FDR corrected permutation t-tests indicating that TL fish swam closer to the center of the tank than all other strains (P’s = 0.0004), and AB spent more time on the periphery (AB compared to TU: P = 0.0006, WIK: P = 0.017). For distance travelled (Figure 2C), we found a main effect of sex (P = 0.0001) where male fish swam further than female fish. There was also a main effect of strain (P = 0.014), but no interaction (P = 0.59). FDR corrected permutation t-tests found that TL fish swam less than AB fish (P = 0.043) and a trend towards a difference between TLs and WIKs (P = 0.061). Finally, for percent of the tank explored (Figure 2D), we also found a main effect of sex (P = 0.024) in which female fish explored less of the tank than their male counterparts. There was also a main effect of strain (P = 0.0001), but no interaction (P = 0.34). Post-hoc tests revealed that TL fish explored the tank less than all other strains (P’s = 0.0004), and that AB fish explored less than TU (P = 0.035) and WIK (P = 0.0026) fish. Taken together, we find that there are several sex differences (center distance, distance travelled, and percent explored) and that TL fish differ the most from other strains across all measures with no clear strain by sex interactions.

**Figure 2.**
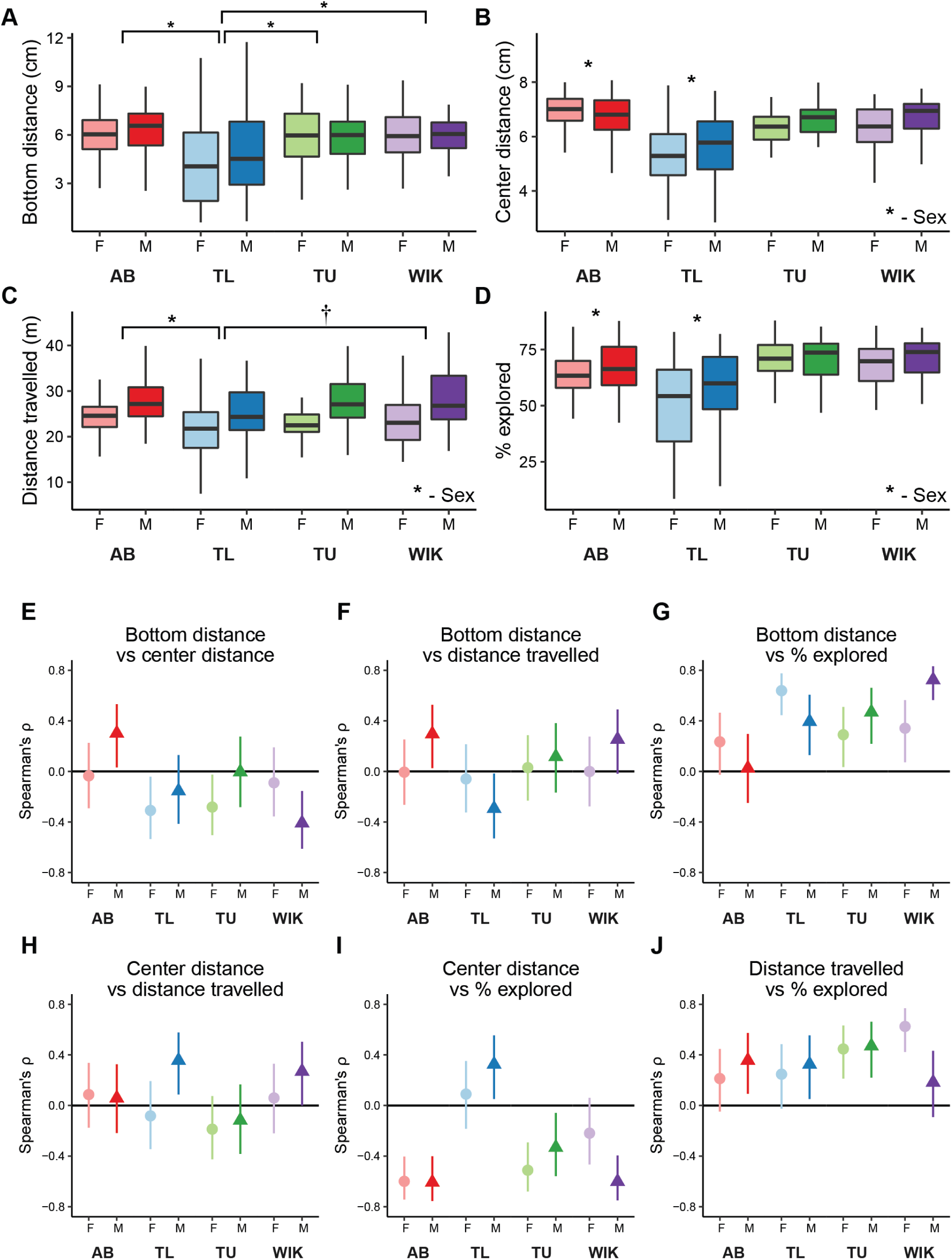
Influence of sex and strain on individual exploratory behaviors and their relationships. The effect of sex and strain on **A**) bottom distance, **B**) center distance, **C**) distance travelled, and **D**) percent of the tank explored. Boxplots indicate median (center line), interquartile range (box ends), and hinge ± 1.5 times the interquartile range (whiskers). Spearman’s rank correlation coefficient (ρ) with 95% confidence intervals across strain and sex for **E**) bottom distance versus center distance, **F**) bottom distance versus distance travelled, **G**) bottom distance versus percent explored, **H**) center distance versus distance travelled, **I**) center distance versus percent explored, and **J**) distance travelled versus percent explored. * - P < 0.05, † - P < 0.10 compared to all other groups or those indicated, n = 50-58.

### Influence of sex and strain on habituation to the novel tank

Some exploratory behaviors have been found to habituate over a single six-minute exposure to a novel tank (Wong et al., 2010), so we examined whether any of the parameters we measured changed over time and were influenced by sex in the various strains (Figure S2). For each measure, we used non-parametric 2 × 6 (sex × time interval) mixed permutation ANOVAs and adjusted for multiple tests using an FDR correction. For distance from bottom (Figure S2A), we found no main effects of sex (AB: P = 0.51, TL: P = 0.46, TU: P = 0.74, WIK: P = 0.86), but found effects of interval in all strains (AB: P = 0.0006, TL: P = 0.0093, TU: P = 0.0006, WIK: P = 0.0084) where all fish, except WIKs, increased their distance from the bottom over time. There were no interactions except for WIKs (AB: P = 0.086, TU: P = 0.74, TL: P = 0.46, WIK: P = 0.018), where female fish appeared to decrease their bottom distance over time whereas male fish showed little change across the trial. For distance from center (Figure S2B), a main effect of interval was found for AB, TU, and WIKs (P’s = 0.0004, TL: P = 0.52), finding that these strains increased their distance from center over time. There were no interactions between sex and time interval (AB: P = 0.91, TL: P = 0.28, TU: P = 0.91, WIK: P = 0.91), but we found trends towards an effect of sex in TU and WIKs (AB: P = 0.28, TL: P = 0.41, TU: P = 0.054, WIK: P = 0.064) where female fish spent their time closer to the center of the tank than male fish, consistent with what was observed in the overall data (Figure 2B). Finally, for distance travelled (Figure S2C), there was an increase in locomotor activity over time in AB, TU, and WIK fish, with a trend in TLs (AB: P = 0.0004, TL: P = 0.093, TU: P = 0.0012, WIK: P = 0.0004). Consistent with the overall data, there were also main effects of sex in all strains except TLs, where there was a trend (AB: P = 0.0014, TL: P = 0.093, TU: P = 0.0004, WIK: P = 0.0048). Only the TU fish had an interaction between time interval and sex where TU female fish increased their distance travelled over time, but male fish did not (AB: P = 0.085, TL: P = 0.39, TU: P = 0.0022, WIK: P = 0.068).

### Correlations between behavioral parameters

To determine the extent to which individual behavioral parameters captured distinct elements of exploratory behavior, and if there was any influence of sex and genetics on these relationships, we computed correlations between individual measures (Figure 2E-J and S3). We used Spearman’s ρ to identify monotonic relationships because of the presence of several non-normally disturbed parameters (Figure S1). As expected, we found that distance travelled was consistently positively correlated with percent explored across all strains and sexes (Figures 2J & S3F). We also found consistent positive correlations between bottom distance and percent explored (Figures 2G & S3C), which is in line with the idea that a higher bottom distance is associated with an increased willingness to explore. However, bottom distance did not correlate consistently with distance travelled (Figures 2F & S3B), suggesting that, despite positive correlations between distance travelled and percent explored, these two exploratory measures are capturing different aspects of exploration. To our surprise, bottom distance and center distance did not consistently correlate with each other (Figures 2E & S3A): in three strain/sexes there was a clear negative correlation (female TLs and TUs, and male WIKs) where fish that swam nearer to the top also swam closer to the center, but in one group (male ABs) the opposite relationship was observed with no clear relationships in the remaining groups. Center distance was mostly negatively correlated with percent explored (Figures 2I & S3E), but not universally so (TLs being the exception), largely consistent with the idea that fish that spend more time closer to the center of the tank also explore more of the tank.

### Identifying behavioral clusters

Given the presence of non-normal behavioral distributions (Figure S1) and variability in the relationship between different exploratory parameters (Figure 2E-J), we hypothesized the presence of multiple behavioral clusters in our data set. To test this, we built a k-nearest neighbor network and applied the Louvain community detection algorithm to identify clusters (Blondel et al., 2008). We used all four behavioral parameters (bottom distance, center distance, distance travelled, and percent explored) in calculating nearest neighbor distances because none of the behavioral measures showed consistently high correlations across all strains and sexes (Figure 2E-J). To determine ‘k’ for building the network, we explored a range of values and chose a value (k = 118) that optimized internal clustering metrics and was robust to small deviations in k (Figure S4) resulting in the identification of four distinct behavioral clusters (Figure 3A).

**Figure 3.**
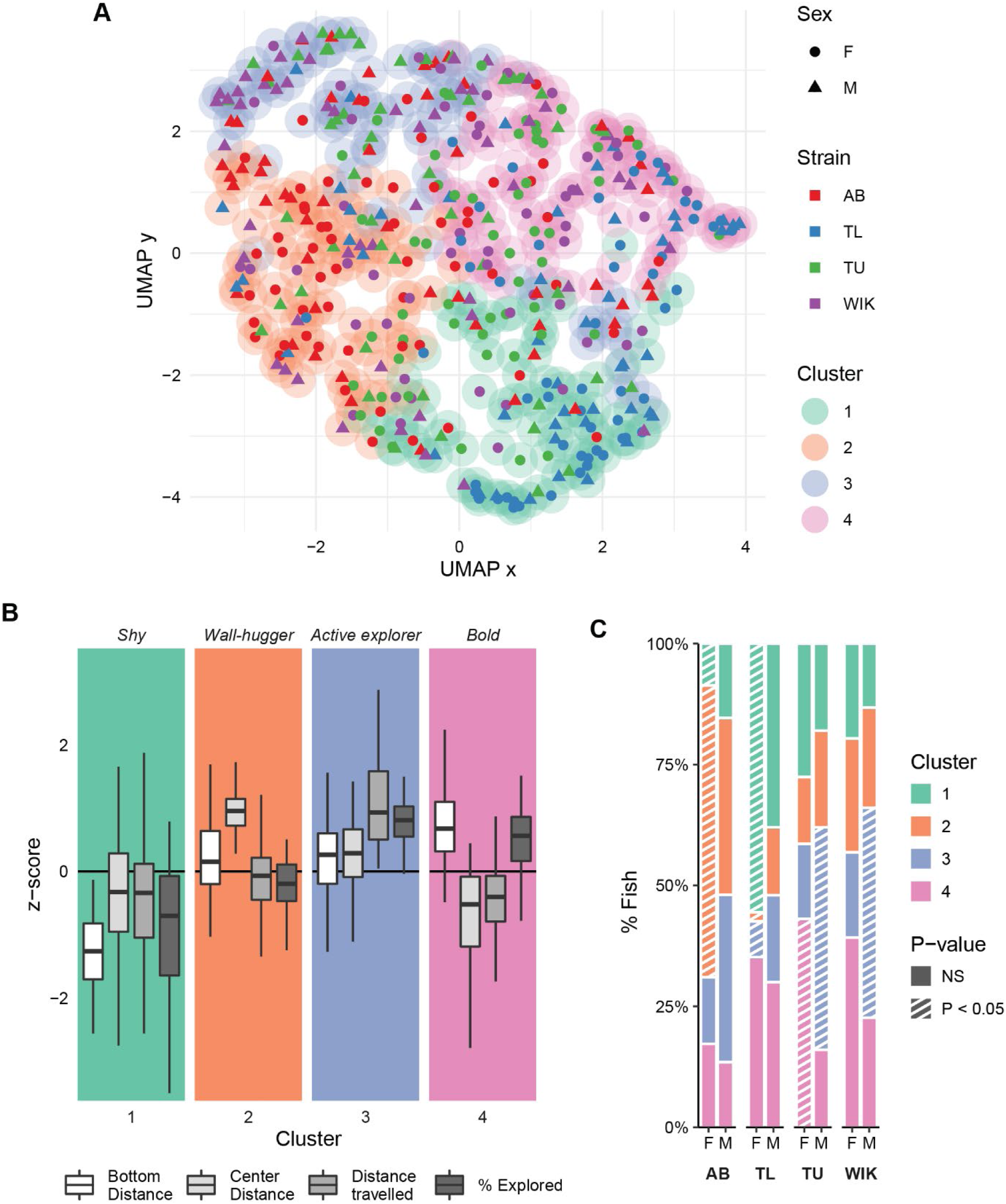
Clustering of zebrafish exploratory behavior during exposure to a novel tank. **A**) Two-dimensional representation of the four-dimensional behavioral space using a uniform manifold approximation (McInnes et al., 2020). Clusters (outer circles) are derived from Louvain community finding applied to a k-nearest neighbor network using 426 fish. Each point is an individual fish where the shape represents the sex (circle: female, triangle: male), inner color the strain, and outer color the behavioral cluster. **B**) Individual behaviors associated with each cluster as box plots indicating median (center line), interquartile range (box ends), and hinge ±1.5 times the interquartile range (whiskers). **C**) Percentage of fish that fall into each cluster across strain and sex. Striped bars: P < 0.05 using randomized permutation tests and FDR corrections, n = 50-58.

An examination of the behaviors associated with each of the clusters reveals a range of behavioral profiles that run the gamut from shy to bold (Figure 3B). Fish in the shyest cluster were lowest in both bottom distance and percent exploration (Figure 3B; cluster 1) whereas fish in the boldest cluster spent the most time near the top and center of the tank while exploring more of the tank than average (Figure 3B; cluster 4). Bold fish were also amongst the lowest in distance travelled, suggesting that they were particularly efficient in their tank exploration. Two ‘mixed’ clusters were also identified: cluster 2 where fish were near average on all measures except for center distance where they spent most of their time near the periphery, a group we call ‘wall-huggers’, and cluster 3 where fish were above average in distance travelled and percent explored that we call ‘active explorers’.

### Behavioral clusters across sex and strain

To determine if fish from a given strain and sex were over- or underrepresented in behavioral clusters, we computed p-values using permutation resampling and FDR corrections for multiple comparisons (Figure 3C). We found that, for the shy cluster (cluster 1), TL fish, irrespective of sex, were overrepresented (female: P = 0.0011, male: P = 0.048), whereas the wall-huggers (cluster 2) had overrepresentation of female AB fish (P = 0.0053). Consistent with our finding that males, on average, swam more than females, more male fish were in the active explorers group (cluster 3), although overrepresentation was only significant in TU and WIK fish (males: AB: P = 0.086, TL: P = 0.23, TU: P = 0.0016, WIK: P = 0.0073). Finally, in the boldest cluster (cluster 4), females outnumbered males in all strains but there was only significant overrepresentation in TU fish with trends in ABs and WIKs (female: AB: P = 0.082, TL: P = 0.16, TU: P = 0.018, WIK: P = 0.064).

### Habituation over two days

All 426 fish used for generating clusters were exposed to the novel tank on two consecutive days, allowing us to determine if their behavior remained consistent across repeated exposures. First, we analyzed individual exploratory behaviors with permutation paired t-tests and FDR corrections (Figure S5). We found that AB and TL fish increased their bottom distance during the second exposure, with a trend in female WIK fish (Figure S5A; female AB: P = 0.0080, male AB: P = 0.012, female TL: P = 0.0016, male TL: P = 0.0040, female TU: P = 0.54, male TU: P = 0.54, female WIK: P = 0.055, male WIK: P = 0.62). Thigmotaxis (center distance) also increased in several groups: AB fish and female WIKs with trends towards an increase in TU females, but a decrease in TL males (Figure S5B; female AB: P = 0.0032, male AB: P = 0.043, female TL: P = 0.38, male TL: P = 0.073, female TU: P = 0.073, male TU: P = 0.82, female WIK: P = 0.043, male WIK: P = 0.87). Distance travelled did not change in any fish (Figure S5C; P’s ≥ 0.12), and percent explored had trends towards a decrease in AB males and increase in TL females (Figure S5D; female AB: P = 0.13, male AB: P = 0.071, female TL: P = 0.071, male TL: P = 0.55, female TU: P = 0.82, male TU: P = 0.39, female WIK: P = 0.55, male WIK: P = 0.31). Taken together, the changes in behavior during the second exposure were mixed, with some changes indicating an increase in behaviors associated with boldness, like bottom distance, and others an increase in putative shy behaviors, like thigmotaxis (however, please see our discussion on thigmotaxis below).

### Behavioral cluster consistency over two days

To determine if the behavioral clusters we identified remained consistent across days, we used exploratory data from the second day of exposure to the novel tank to assign fish to clusters uncovered on the first day (Figure 4A). We found that 52.8% of fish fell into the same behavioral cluster on days 1 and 2 (P = 0.0001; permutation average = 25.1% overlap). Overall, the number of fish falling into the shyest group (cluster 1) decreased, and the wall-huggers (cluster 2) remaining the most consistent. At the level of sex and strain (Figure 4B), we again used FDR corrected permutation resampling, finding that, as on day 1, AB females were overrepresented in the wall-huggers group (cluster 2; P = 0.0011), but were now joined by their male counterparts (P = 0.031). TL females also remained overrepresented in the shyest cluster (P = 0.0011), but a significant portion of male and female TL fish were now in the boldest cluster (female TL: P = 0.013, male TL: P = 0.0011). TU females were no longer overrepresented in the boldest group (P = 0.62), although TU males were now underrepresented in this group (P = 0.025) and maintained their overrepresentation as active explorers, cluster 3 (P = 0.0011). Finally, neither female nor male WIKs were overrepresented in any cluster on day 2.

**Figure 4.**
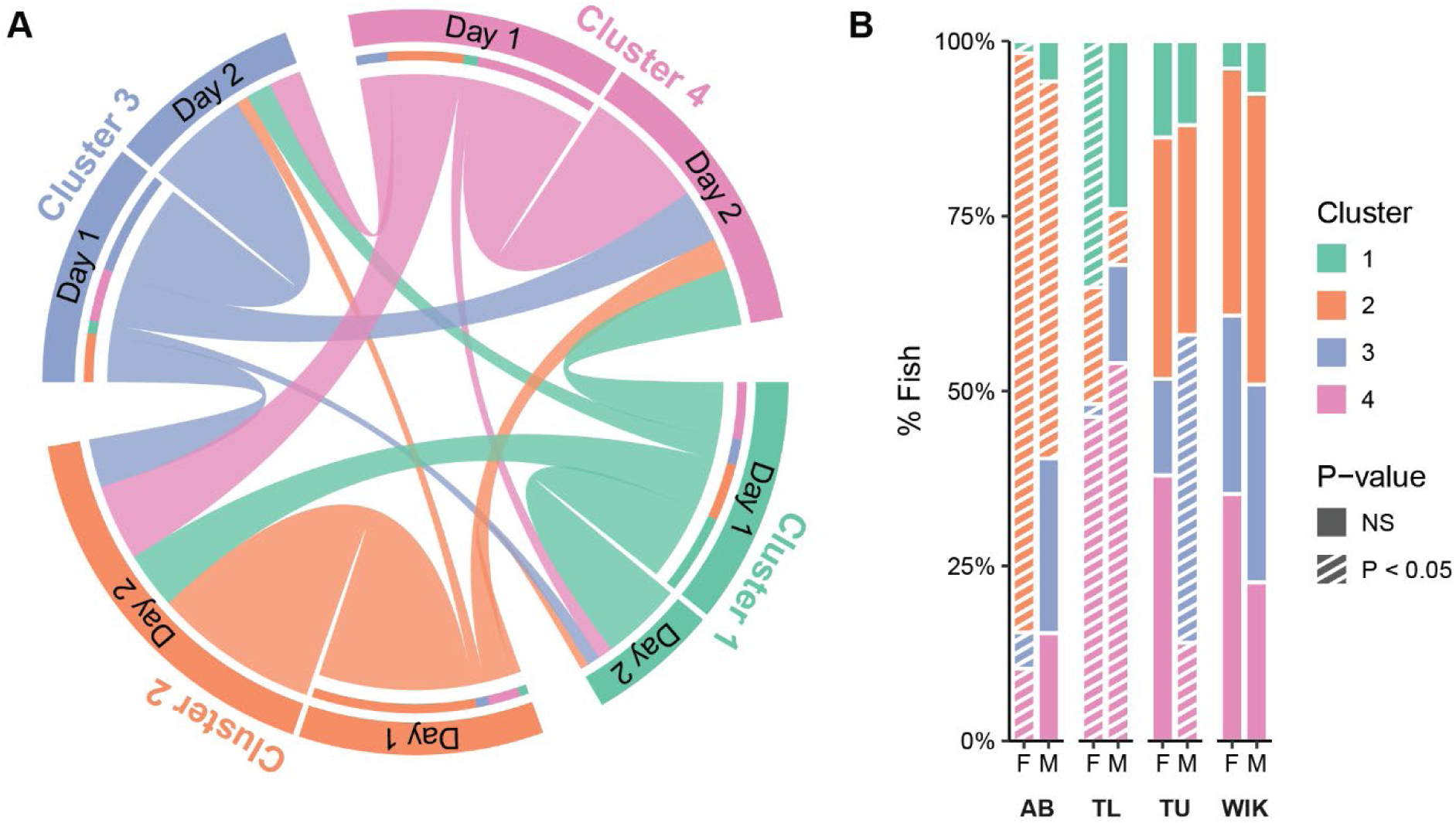
Clusters across two consecutive exposures to the novel tank. **A**) Chord diagram indicating how cluster membership changes from day 1 to day 2 of novel tank exposure. **B**) Percentage of fish that fall into each cluster across strain and sex on the second day of exposure to the novel tank. Striped bars: P < 0.05 using randomized permutation tests and FDR corrections, n = 50-58.

### Habituation over multiple days and weeks

Some exploratory behaviors have been found to habituate over several days of exposure to a novel tank (Wong et al., 2010). In a new cohort of AB and TU fish, exploratory behavior was measured over five consecutive days of exposure to the novel tank (Figure S6). We used 5 × 2 (day × sex) mixed permutation ANOVAs to assess significance. For bottom distance in AB fish (Figure S6A), there was no effect of day (P = 0.20), sex (P = 0.14) or an interaction (P = 0.64), whereas TU fish increased their bottom distance over the five days (P = 0.025), with no effect of sex (P = 0.18) nor an interaction (P = 0.87). AB fish increased their center distance (Figure S6B) over time (P = 0.0001) with no effect of sex (P = 0.12) or an interaction (P = 0.14), and TU fish had a similar trend (day: P = 0.094, sex: P = 0.27, interaction: P = 0.93). As we saw before, male AB fish swam further than female fish (P = 0.010) with no effect of day (P = 0.62). There was a trend towards and interaction between day and sex (P = 0.058) where AB males swam less over time, and females more (Figure S6C). In TU fish, there was a trend towards males swimming more than females (P = 0.084), and a clear effect of day (P = 0.0007) and no interaction (P = 0.68) as both sexes increased their distance travelled over time. Finally, for percent explored (Figure S6D), In AB fish there was no effect of sex (P = 0.33) or day (P = 0.77), but there was a trend towards an interaction (P 0.068) where female fish appeared to slightly decrease their percent explored over time. In TU fish, animals increased their exploration over days (P = 0.0089), with a trend towards males exploring more than females (P = 0.099), and no interaction of day and sex (P = 0.31).

In a separate cohort of TU fish, we examined individual exploratory behaviors over 10 weeks of biweekly (every other week) exposures to the novel tank (Figure S7). We used 6 × 2 (week × sex) mixed permutation ANOVAs to assess significance. For bottom distance (Figure S7A), we found no effect of week (P = 0.96) or sex (P = 0.23) but there was an interaction (P = 0.0023) where female fish increased, and males decreased, their distance from bottom across weeks. For center distance (Figure S7B), there was a trend towards an effect of week (P = 0.076), and no effect of sex (P = 0.49) nor an interaction (P = 0.88). For both distance travelled (Figure S7C) and percent explored (Figure S7D), there were main effects of sex (P = 0.0001 and P = 0.015, respectively) where, as we saw before, male fish swam further, and explored more of the tank, than female fish. There were also main effects of week (P = 0.0001 and P = 0.0002, respectively) and no interactions (P = 0.80 and P = 0.25, respectively), where both female and male fish increased their exploratory behaviors during repeated exposures.

### Behavioral cluster consistency over days and weeks

Next, we asked whether the behavioral clusters of individual animals over five consecutive days or across 10 weeks remained consistent (Figure 5). Across the exposures we found that exploratory behavior of approximately 50% of animals fell into the same cluster on at least 4 out of 5 days (Figure 5A) or 5 of 6 biweekly exposures (Figure 5B). To determine if the consistency across time was greater than chance, each animal was assigned an overlap score: the sum of pair-wise overlaps across consecutive exposures to the tank. For the daily data this score ranged from 1 (only one pair of days overlapped) to 10 (all pair-wise overlaps), for the biweekly data it ranged from 2 to 15. During 5 days of exposure the average scores for AB and TU fish were 5.6 ± 2.6 and 5.9 ± 3.1 (mean ± standard deviation), respectively, both of which were significantly higher than the overlap scores from resampling (permutation mean ± standard deviation: 3.4 ± 0.2 and 2.7 ± 0.2, respectively; P’s = 0.0002). For the biweekly data, the average overlap score was 9.0 ± 4.2, which was significantly higher than chance (4.8 ± 0.2, P = 0.0002). Although most animals had scores higher than the permutation average, a small subset of animals were “consistently inconsistent” with their exploratory behavior falling into all four clusters across days or weeks. In both the daily and biweekly data, we found that the number of animals falling into the shyest cluster (cluster 1) decreased over time (Figure S8). There were few prominent changes in other clusters except for an increase in wall-huggers (cluster 2) in AB fish over five consecutive days (Figure S8A) and an increase in active explorers (cluster 3) during biweekly exposures (Figure S8B).

**Figure 5.**
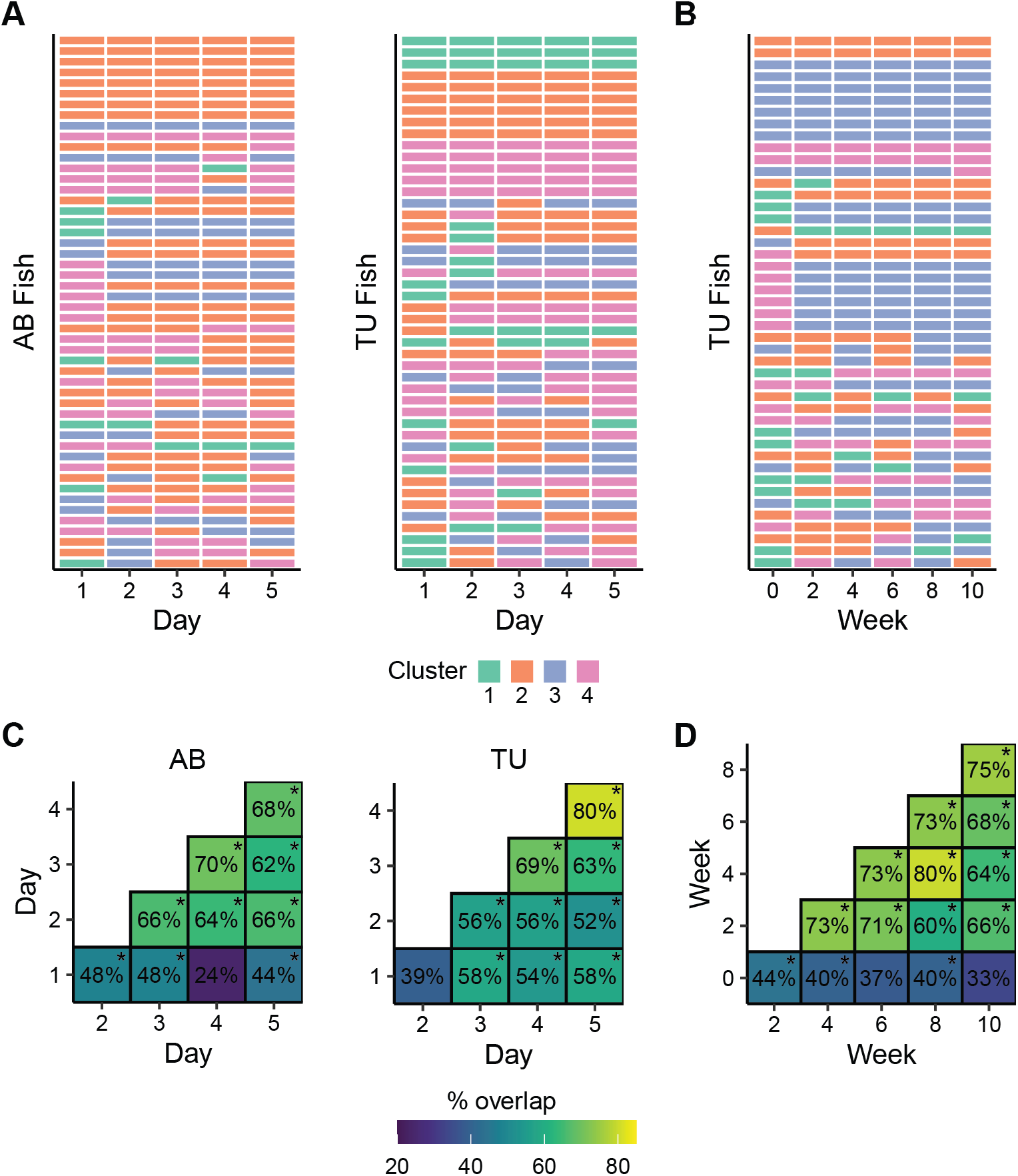
Behavioral consistency in the novel tank across days and weeks. **A**) Cluster membership of AB (n = 46) or TU (n = 50) fish that were exposed to the novel tank on five consecutive days. **B**) Cluster membership of TU (n = 45) fish that were exposed to the novel tank every other week for 10 weeks. **C**) Percent of overlapping clusters in AB and TU fish across consecutive exposures to the novel tank across five days. **D**) Percent overlapping clusters in TU fish across biweekly (every other week) exposures to the novel tank. * - P < 0.05 using randomized permutation test and FDR corrections.

Finally, we examined how much cluster overlap there was between each of the five days, or weeks, of the experiment (Figure 5C and D). During the daily exposures, we found that the clusters from day 1 overlapped above chance with most clusters on subsequent days in both AB and TU fish, with the overlap in TU fish averaging about 10% higher than AB fish (Figure 5C). In AB fish, cluster overlap after day 2 increased markedly (range 62-70%). In TU fish, an increase in overlapping clusters did not occur until day 3, where it also remained high (63-80%). We saw a similar trend during biweekly exposures (Figure 5D) where cluster identity on the first day overlapped the least with subsequent weeks (33-44%), but from the second to the tenth week, overlap was considerably higher (60-80%).

## Discussion

By applying an unbiased approach to the clustering of three-dimensional exploratory behavior from over 400 zebrafish, we found that behavior stratifies into four distinct behavioral clusters. These profiles included previously described bold and shy behaviors as well as two novel behavioral types we call wall-huggers and active explorers. Notably, these individual differences in fish behavior were consistent over days and weeks, one of the key hallmarks of personality. Although there were few strain-sex interactions on individual behaviors, the distribution of clusters varied considerably across strain and sex suggesting biological modulation of these behavioral clusters.

Studies that have examined individual differences in exploratory behavior typically assume a bimodal distribution of bold versus shy. Here, enabled by our large dataset of three-dimensional behavior, we tested this assumption. Using an unbiased unsupervised machine learning approach, we found four distinct behavioral subtypes (shy, wall-huggers, active explorers, and bold). Cluster 4 was the boldest cluster where fish behavior corresponded to traditional descriptions of boldness: above average time in the top and center of the tank and greater exploration. The behavior of fish in cluster 1 was designated as shy because these fish spent most of their time near the bottom of the tank. Interestingly, in this shy group, behaviors like center distance and percent explored exhibited a high degree of variability, suggesting the potential presence of additional subgroups that would require an even larger data set to uncover.

Of the novel behavioral clusters we uncovered, we designated cluster 2 ‘wall-huggers’ because these fish spent most of their time towards the periphery of the tank and were near average on all other parameters. We do not refer to this group as ‘shy’ because, although thigmotaxis has been interpreted as a predator avoidance or anxiety-like behavior in zebrafish (Kalueff et al., 2013), our findings do not clearly support this interpretation. For example, we did not observe consistent negative correlations between center and bottom distance (Figure 2E), as would be expected if these behaviors reflected the same underlying construct (i.e. predator avoidance). We also find that, unlike bottom dwelling, thigmotaxis increased over time within and between sessions of the novel tank test (Figures S2B and S6B), the opposite of the habituation we would expect to see if thigmotaxis was an anxiety-like or predator avoidance behavior. Our findings are consistent with other studies that examined thigmotaxis over time and/or alongside other avoidance behaviors in adult fish (Blaser et al., 2010; Champagne et al., 2010; Rosa et al., 2018; Shams et al., 2015; Wong et al., 2012). It may be that the interpretation of thigmotaxis in zebrafish has been unduly influenced by findings in rodents where its importance for predator avoidance is clearer (Champagne et al., 2010; Simon et al., 1994). Thus, we propose that thigmotaxis in adult zebrafish should not be interpreted in the context of predator avoidance or anxiety until its relevance for fish can be clarified.

We designated behavioral cluster 3 as ‘active explorers’ because these fish were clearly above average in both distance travelled and percent explored, but near average on center and bottom distance. This group of fish may correspond to what has been referred to as low stationary, or proactive, zebrafish in other work (Baker et al., 2018; Wong et al., 2012). In comparison to the boldest cluster, active explorers are distinguished by a notable elevation in distance travelled with similar levels of percent exploration. Thus, during exploration, these animals are less efficient in their exploration as they revisit many parts of the tank. This suggests that these animals may have poor working memory or higher resting metabolic rates, the latter having been found to be predictive of individual differences in activity (Biro and Stamps, 2010; Yuan et al., 2018).

Strain and sex had a significant influence on behavioral cluster identity. During the initial exposure to the tank, female fish more often fell into the boldest cluster, although only TU females were statistically overrepresented. Male fish were more likely to be in the active explorer group, particularly TU and WIK male fish. This likely reflects the fact that, on average, male fish tended to swim further, and explore more of the tank, than female fish. This increase in locomotor activity in male fish was expected as it has been reported elsewhere (Ariyomo and Watt, 2015; Clayman et al., 2017; Philpott et al., 2012), although not universally so (Ampatzis and Dermon, 2016; Fontana et al., 2019; Rambo et al., 2017; Tran and Gerlai, 2013). We also found interactions between sex and strain on cluster identity, such as underrepresentation of AB females, and overrepresentation of TL females, in the shyest cluster with the situation reversed in the wall-huggers (cluster 2) group; notably, this interaction persisted into the second day of testing, suggesting it is particularly robust.

The behavioral clusters we identified demonstrated a high degree of consistency across days and weeks, supporting the idea that they are akin to personality types (Gosling, 2001). In our initial experiment, where fish were tested in the novel tank on two consecutive days, overlap was above 50%, twice as high as would be expected by chance (∼25%). Upon repeated exposures to the tank over five consecutive days we found that cluster overlap steadily increased to above 65%, and over half of the fish were in the same cluster on 4 of the 5 days.

We found similar effects during biweekly (every other week) exposures over 10 weeks. This increase in overlap over days is likely due, in part, to habituation (Wong et al., 2010) given that we found that there were main effects of day on several individual measures, particularly in TU fish (Figures S6 and S7). Nonetheless, this is the first report, to our knowledge, of consistency in zebrafish behavior across days and weeks using unbiased multidimensional behavioral clustering that approximates key elements of personality.

The present work also yielded a comprehensive account of how genetic background influences individual exploratory behaviors of zebrafish, adding to an already rich, if inconsistent, literature. The primary strain difference on individual behaviors we observed was that TL fish differed from most other strains, demonstrating higher bottom dwelling, less thigmotaxis, and exploring less of the tank. Although strain differences in zebrafish behavior during the novel tank test have been reported (Egan et al., 2009; Maximino et al., 2013; Mustafa et al., 2019), only a few studies have compared widely used inbred strains. Vignet and colleagues (2013) found that AB fish exhibited more bottom dwelling than TU fish, and Audira et al (2020) found no differences between AB, TL, and WIK fish in bottom dwelling or distance travelled, but that WIK fish exhibited greater thigmotaxis. Others have found AB fish to exhibit greater bottom dwelling than WIKs (Sackerman et al., 2010), or a decrease in bottom dwelling in WIK fish over 60 minutes, but not TU fish, with no difference in locomotor activity (Pannia et al., 2014). Differences between these studies, and the present work, may be attributable to unreported sex ratios, housing conditions, nuances of the behavioral task, or genetic drift such that inbred strains from different labs or suppliers may differ subtly, as has been observed in rodents (Taft et al., 2006). We attempted to address some of these challenges in our study. For example, housing conditions are known to influence zebrafish behavior in the novel tank test (Parker et al., 2012; Reolon et al., 2018). We housed all animals in mixed sex pairs for one week prior to behavioral testing. This allowed us to maintain zebrafish identify over time while avoiding any effects of stress due to social isolation or tagging. To minimize genetic drift, and maximize the likelihood our findings will translate to other labs, all fish were within 2 generations of breeders obtained from the Zebrafish International Resource Center where they maintain a much larger, and genetically diverse, population of animals. Nonetheless, given that inbred zebrafish lines are not isogenic (Nasiadka and Clark, 2012), there is no obvious way to ensure genetic similarity of lines across labs.

Taken together, we have found that the exploratory behavior of zebrafish goes beyond bold versus shy, stratifying instead into four distinct categories. This finding was enabled by recent advances in animal tracking and 3D video capture that allowed us to scale up our behavioral assessment using inexpensive open-source tools. As would be expected of behaviors capturing personality, the clusters we identified are consistent over days and weeks and influenced by strain and sex. Future work will be needed to determine if these clusters are predictive of behaviors in other contexts. Nonetheless, our findings suggests that zebrafish behavior is more complex than is typically assumed and that this complexity needs to be taken into consideration when using zebrafish to study the biological basis for individual differences in behavior.

## Acknowledgments

We thank Barbara D. Fontana for excellent comments on a prior version of this manuscript. This work was funded by the National Institutes of Health (R35GM142566) to JWK.

## Methods

### Subjects

Subjects were female and male AB, TU, WIK, or TL zebrafish 16-32 weeks of age. All fish used in experiments were bred and raised at Wayne State University and within two generations of animals obtained from the Zebrafish International Resource Center at the University of Oregon. Animals were kept on high density racks under standard conditions (temperature 26.5 ± 0.5 C; water conductivity 500 ± 10 μS, and a pH of 7.5 ± 0.2) with a 14:10 light/dark cycle (lights on at 8:00AM). Fish were fed twice a day with a dry feed in the morning and brine shrimp (*Artemia salina*; Brine Shrimp Direct, Ogden, UT) in the afternoon. Behavioral testing took place between 11:00 and 14:00. Sex of fish was confirmed via dissection after behavioral procedures were complete. Those animals that were assigned the wrong sex were removed from analysis. All procedures were approved by the Wayne State University Institutional Animal Care and Use Committee.

### Behavioral apparatus

Five-sided novel tanks (15 x 15 x 15 cm) were made from frosted acrylic (ShopPopDisplays, Woodland Park, NJ) and open from above. Tanks were placed in an enclosure made of white plasticore to diffuse light and prevent fish from seeing experimenters. Intel RealSense^TM^ cameras (D435) were mounted 20 cm above the tank to capture three-dimensional videos (Kuroda, 2018). Video capture was done using custom written Python scripts and the pyrealsense2 package.

### Behavioral procedures

One week prior to behavioral testing, fish were placed as male/female pairs into 2 L tanks. The tanks were divided in half with a transparent divider with two fish in each section and a total of four fish in each tank. This allowed us to maintain the identity of fish over days without isolation while also creating a consistent social environment across all animals. On days when behavior was assessed, fish were taken off housing racks and moved to the procedural space at least one hour prior to behavioral testing. Following testing, fish sat for one hour before being returned to the housing racks. The novel tanks were filled with 2.5 L of fish facility water and individual fish were placed in the tanks for six minutes while video was recorded for offline analysis. Novel tanks were rinsed between animals and water was replaced.

### Animal Tracking

Fish were tracked in the color videos using DeepLabCut (Mathis et al., 2018). We tracked five points (head, trunk, and three points on the tail; Figure 1C). Using ResNet 101, we initially trained the network on 160 frames equally divided across fish of all four strains and both sexes. We refined and improved our initial training by correcting outliers and including an additional 160 frames.

To obtain the z-coordinate of fish at each frame, we overlaid the tracked points at each color frame with the depth stream from the cameras. One z-coordinate for the fish was identified for each frame based on a 4-pixel search radius around the tracked points starting with the trunk. If no point was identified from the depth stream, it was interpolated. Z-coordinates were corrected for the diffraction of water by measuring 100 points of varying distance from the camera in the presence and absence of water and using the slope of a least-squares fit line.

### Exploratory Behavioral Parameters

To measure bottom distance, we calculated the equation for a plane along the bottom of the tank using least squares fit of four points. We then calculated the shortest distance between a point (the fish) and the plane. To measure center distance, we calculated how far fish were from a line made from points at the center top and bottom of the tank. Percent of the tank explored was calculated by dividing the tank into 1,000 evenly spaced voxels and calculating the number of unique voxels visited. For distance travelled, traces were first smoothed using a Savitzky-Golay filter with a length of 7 frames and an order of 3 (Press and Teukolsky, 1990) and then Euclidean distance was calculated between points of successive frames.

### Behavioral clusters

We identified behavioral clusters using a Louvain community detection algorithm (Blondel et al., 2008) applied to a k-nearest neighbor network on our initial dataset of 426 fish during their first day of exposure to the novel tank. This was done by first standardizing individual behavioral parameters (bottom distance, center distance, distance travelled, and percent explored), and calculating a similarity score between each fish:

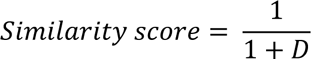

Where D is the Euclidean distance between each fish in four-dimensional behavioral space. To determine the best k for building the network, we initially used a range of k’s, applied the Louvain community finding algorithm to weighted, non-directed networks, and calculated internal clustering metrics (Calinski-Harabsz index (Caliński and Harabasz, 1974), Silhouette index (Rousseeuw, 1987), and Davies-Bouldin index (Davies and Bouldin, 1979)). We chose k=118, which was in the middle of a regime that optimized internal clustering and was robust to small changes in k (Figure S4).

To identify clusters in new data (e.g. the second day of novel tank exposure (Figure 4) fish exposed on consecutive days or biweekly (Figure 5)), we first standardized the new data using the parameters from the 426 fish exposed to the novel tank described above. Then, for each new data point, we assigned clusters based on the proportion of connections to its 51 nearest neighbors in the initial network where 51 is half the size of the smallest cluster identified.

### Coding and statistical analysis

Statistical analysis was performed using R version 4.1.2 (R Core Team, 2016) and visualized using ggplot2 (Wickham, 2015) and RColorBrewer (Neuwirth and Brewer, 2014). Normality was assessed using the Shapiro-Wilks test. Because a considerable portion of the data was not normally distributed, we used permutation ANOVAs using the permuco package (Frossard and Renaud, 2021), and the RVAideMemoire (Hervé and Hervé, 2020) for permutation t-tests. Multiple comparisons were corrected using a false discover rate (FDR) correction (Benjamini and Hochberg, 1995). We used packages cccd (Marchette et al., 2015) and igraph (Csardi and Nepusz, 2006) to build and analyze the k-nearest neighbor network, and the ClusterCrit (Desgraupes, 2013) to assess internal clustering metrics. The chord diagram was made using circlize (Gu et al., 2014).

For all permutation tests, we resampled data 10,000 times without replacement and calculated the P-value as the proportion of experimental observations that were more extreme than the permutation observations. When multiple tests were performed, P-values were corrected using the FDR as indicated. To determine if behavioral clusters were affected by strain and sex (Figure 3C and 4B), we resampled cluster assignment and calculated how many animals from each strain and sex fell into each cluster. For comparison of behavioral clusters across two days (Figure 4B) we resampled cluster assignments from day 2. To determine if behavior was consistent across daily (five day) or biweekly (every other week over ten weeks) (Figure 5A-B), we first calculated an overlap score for each fish that was the sum of the number of days or weeks that had overlapping clusters (ranging from 1 to 10 for five-day data, 2 to 15 for biweekly data). We then took the average of these overlap scores and compared them to scores obtained from permutation resampling (without replacement) of cluster assignments.

Five-day and biweekly percent overlap calculations and permutation tests were as described for the two-day data except that resampling of cluster assignments was done individually for all days/weeks except the fist day/week.

## Supplemental Information

**Figure S1.**
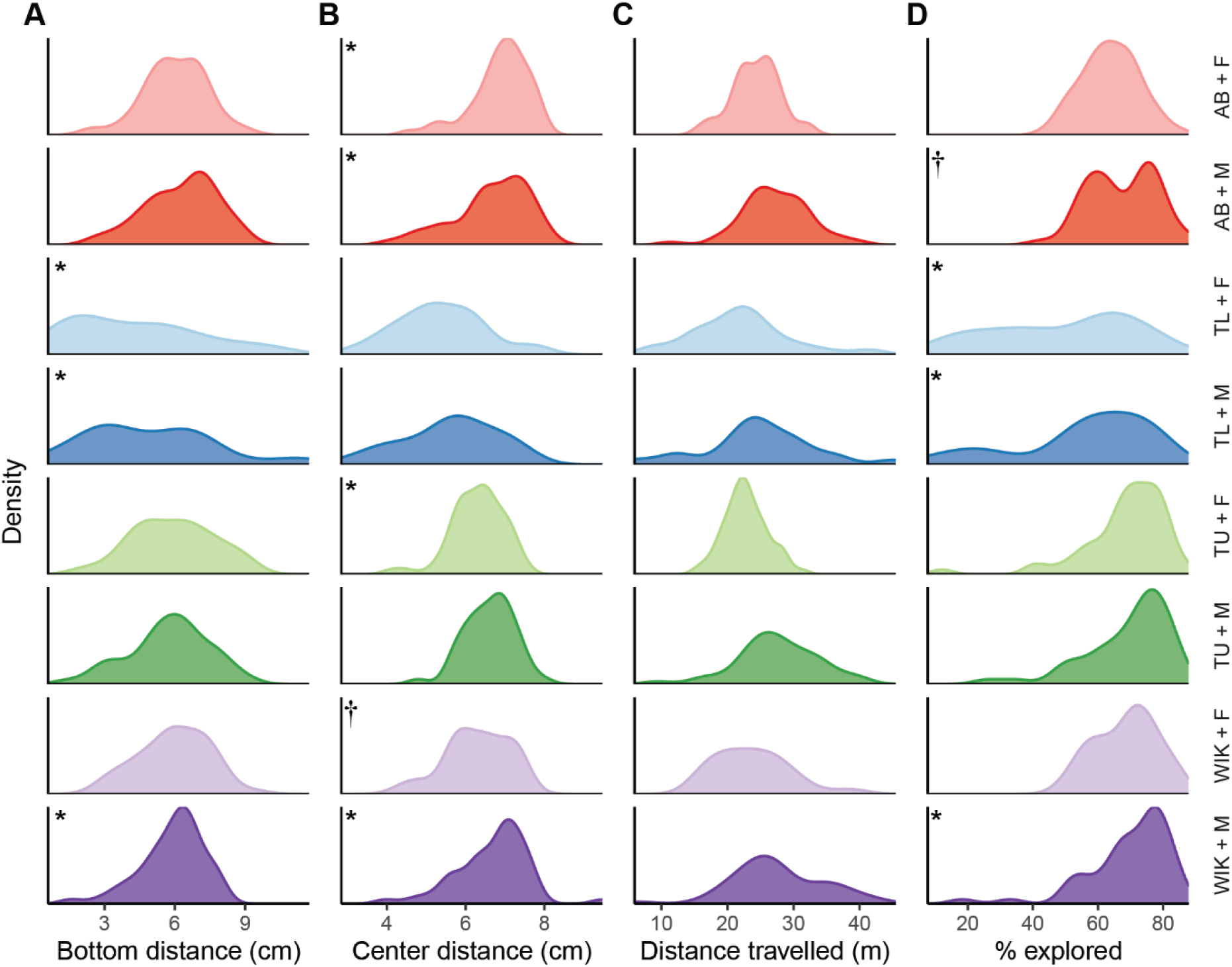
Distribution of values for individual behavioral parameters across stain and sex. Distributions for **A**) bottom distance, **B**) center distance, **C**) distance travelled, and **D**) percent tank explored. * - P < 0.05, † - P < 0.10 Shapiro-Wilks test for normality, n = 50-58.

**Figure S2.**
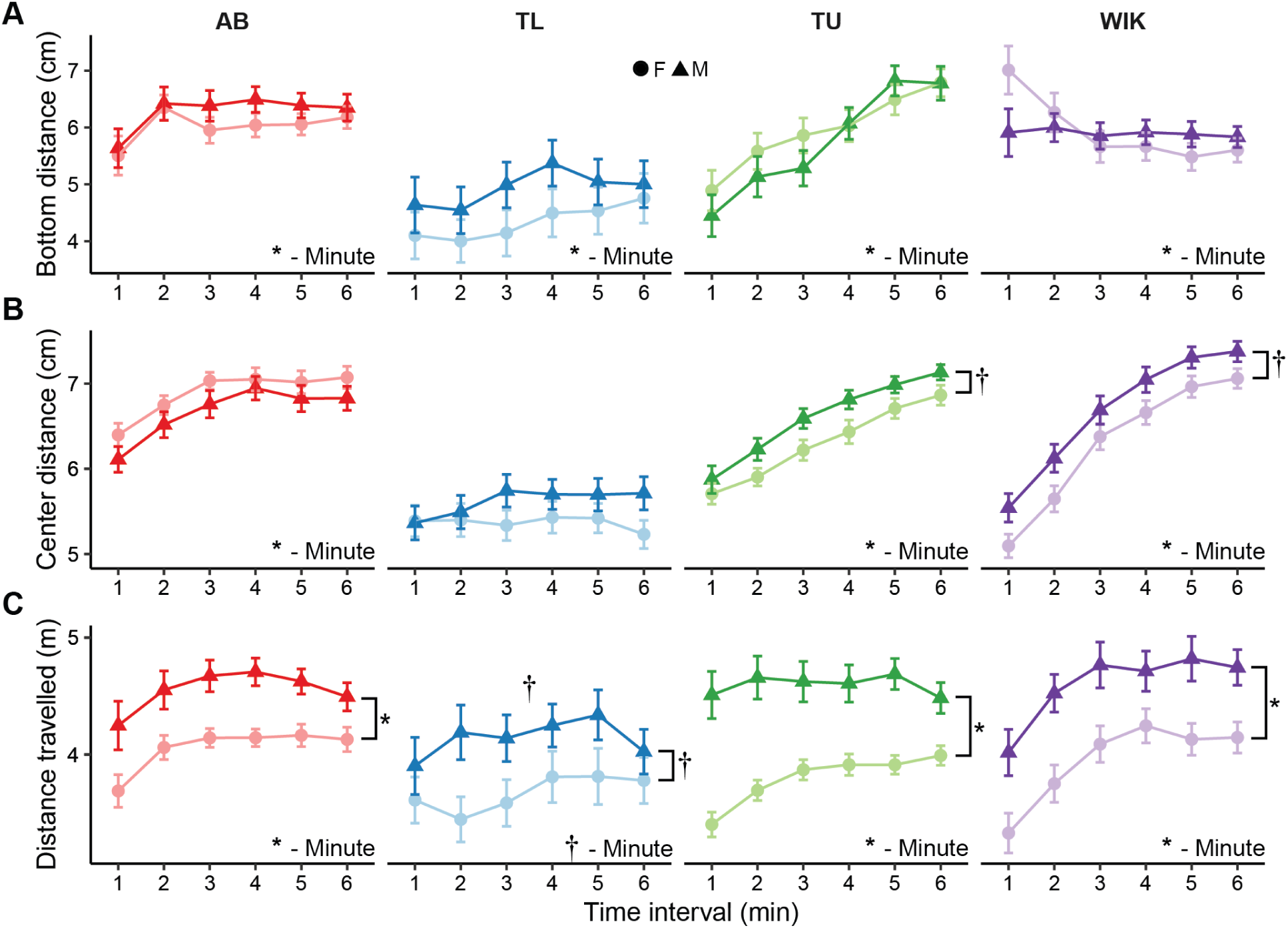
Individual exploratory behaviors across six minutes during a single exposure to the novel tank. The effect of strain (color) and sex (shape, circle: female, triangle: male) across time for **A**) bottom distance, **B**) center distance, and **C**) distance travelled. * - P < 0.05, † - P < 0.10 for time interval (minute) or sex as indicated, n = 50-58. Data presented as mean ± SEM.

**Figure S3.**
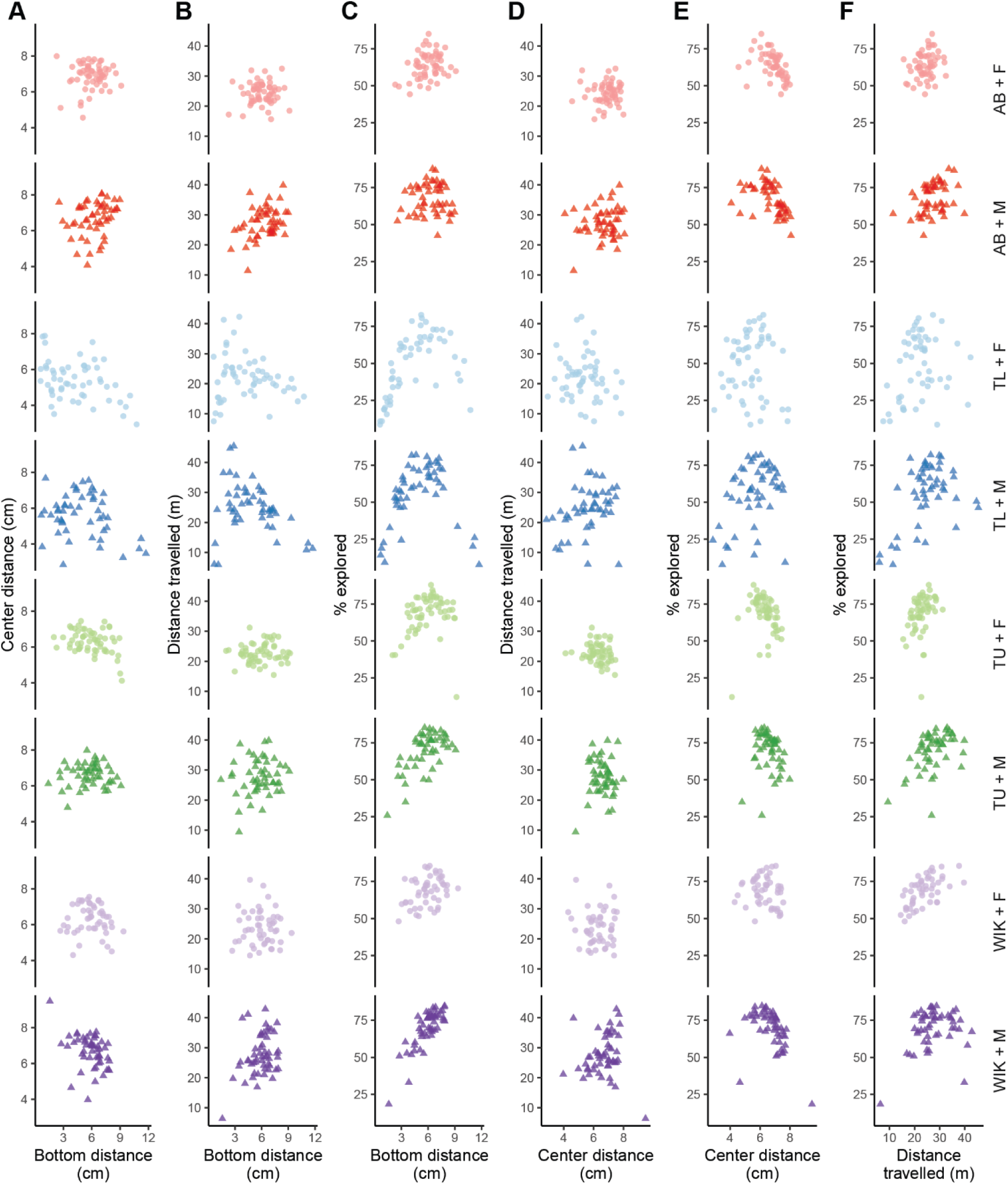
Relationship between pair-wise individual behavioral parameters. Scatterplots across strain and sex of **A**) bottom distance versus center distance, **B**) bottom distance versus distance travelled, **C**) bottom distance versus percent explored, **D**) center distance versus distance travelled, **E**) center distance versus percent explored, and **F**) distance travelled versus percent explored. n = 50-58.

**Figure S4.**
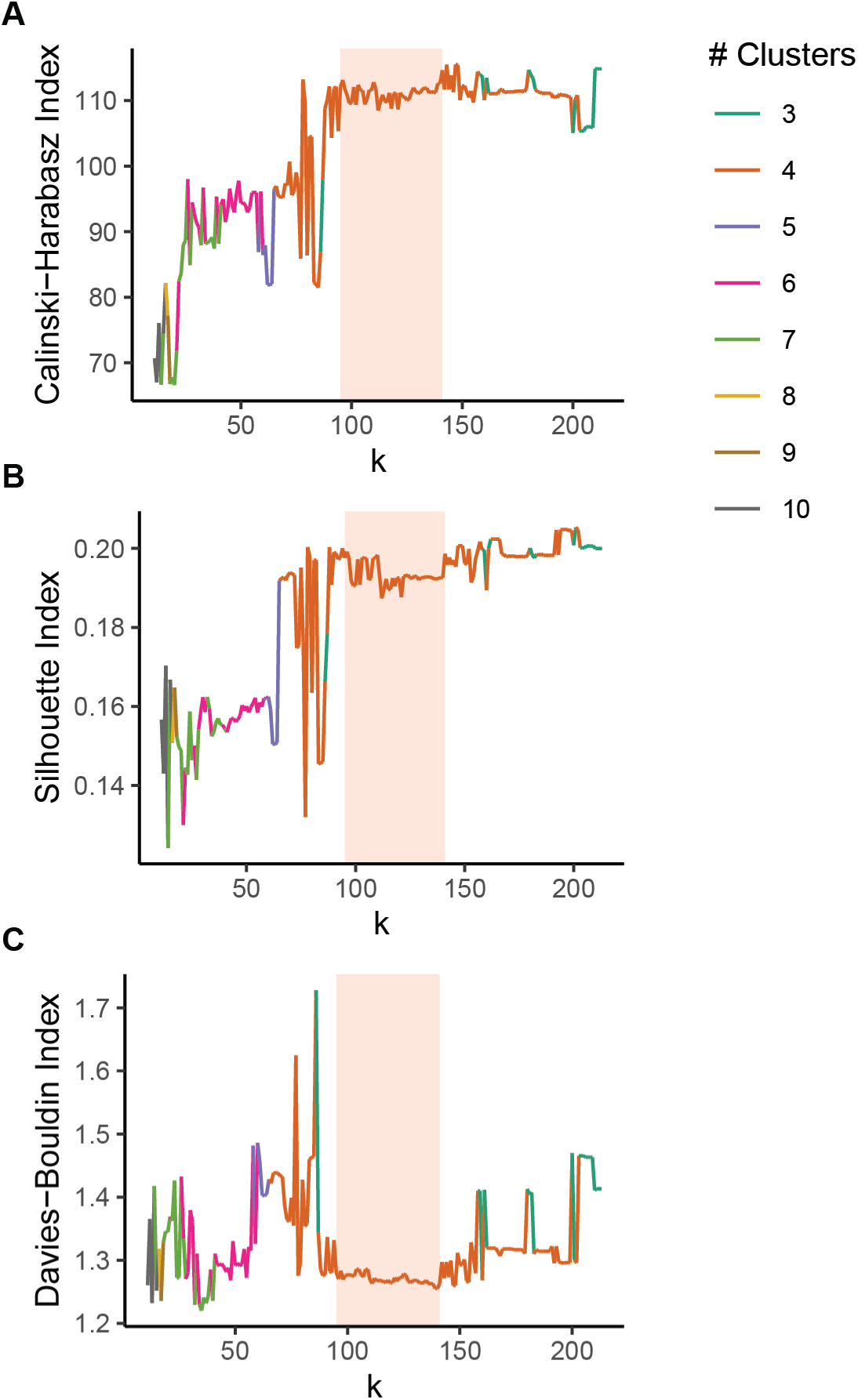
Internal clustering metrics for knn Louvain clustering. The parameter k was varied for generating knn’s and Louvain clustering followed by measuring the **A**) Calinski-Harbasz, **B**) silhouette, and **C**) Davies-Bouldin indices at each k. Area highlighted in red indicates region where clustering metrics were near optimal and robust to small changes in k.

**Figure S5.**
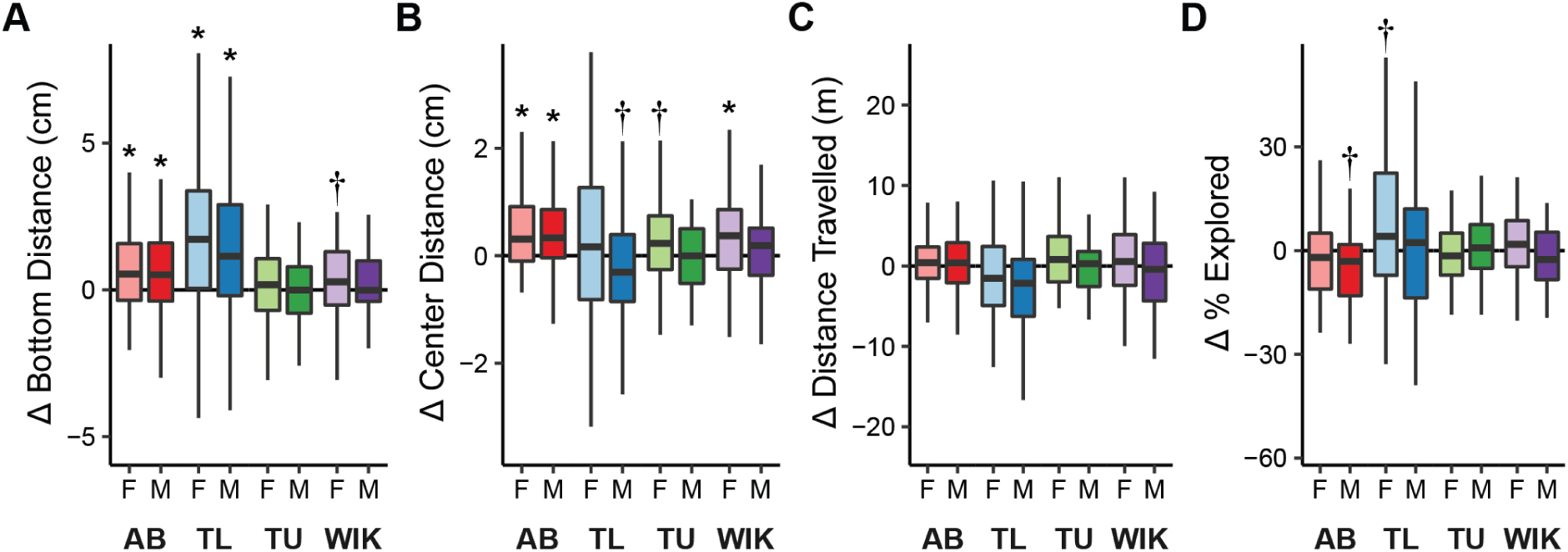
Changes in individual exploratory behaviors during two consecutive exposures to the novel tank. The effect of strain and sex on changes (day 2 minus day 1) over two days in **A**) bottom distance, **B**) center distance, **C**) distance travelled, and **D**) percent explored. Boxplots indicate median (center line), interquartile range (box ends), and hinge ± 1.5 times the interquartile range (whiskers). * - P < 0.05, † - P < 0.10 compared to zero, n = 50-58.

**Figure S6.**
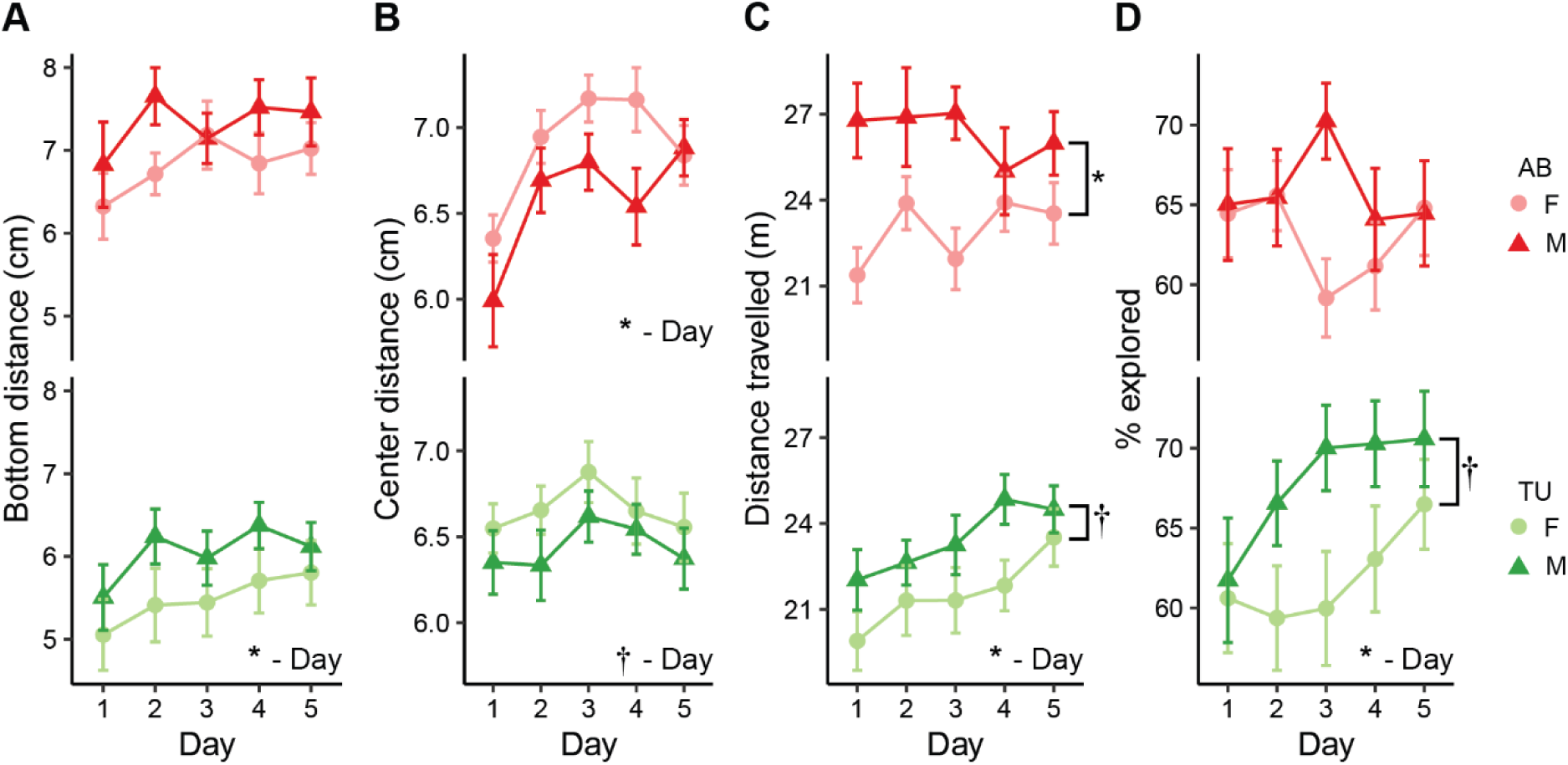
Individual exploratory behaviors during five consecutive days of exposure to the novel tank. AB (top) and TU (bottom) fish were exposed to the novel tank on five consecutive days and **A**) bottom distance, **B**) center distance, **C**) distance travelled, and **D**) percent explored were measured. * - P < 0.05, † - P < 0.10 for day or sex and indicated, n = 21-26. Data presented as mean ± SEM.

**Figure S7.**
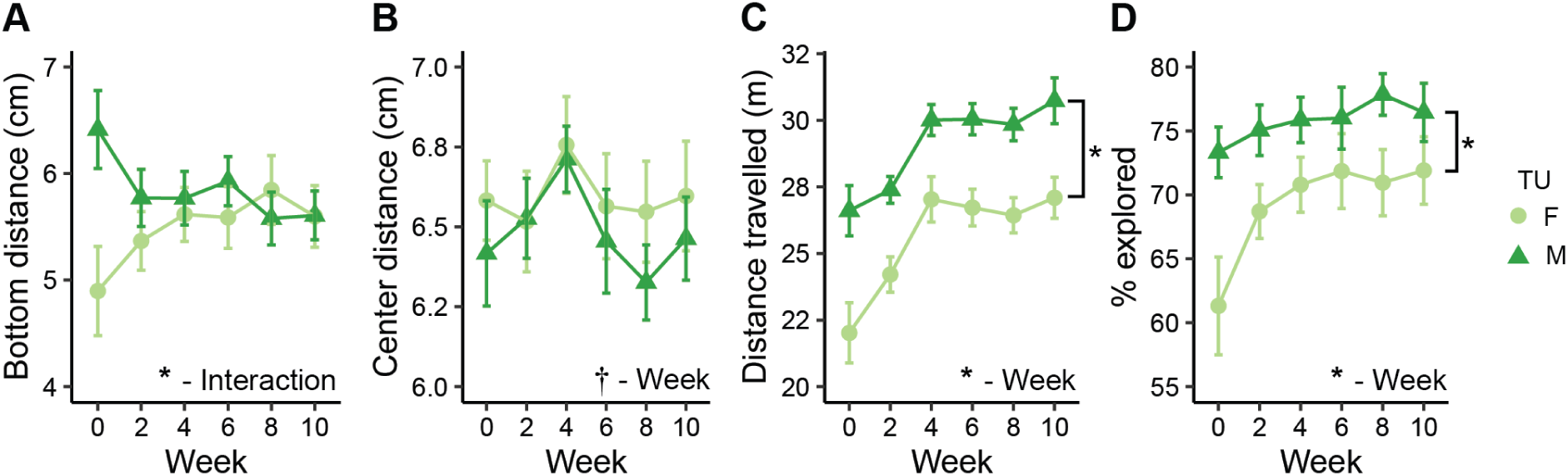
Individual exploratory behaviors over ten weeks. TU fish were exposed to the novel tank every other week and we measured **A**) bottom distance, **B**) center distance, **C**) distance travelled, and **D**) percent explored. * - P < 0.05, † - P < 0.10 for week or sex as indicated, n = 22-23. Data presented as mean ± SEM.

**Figure S8.**
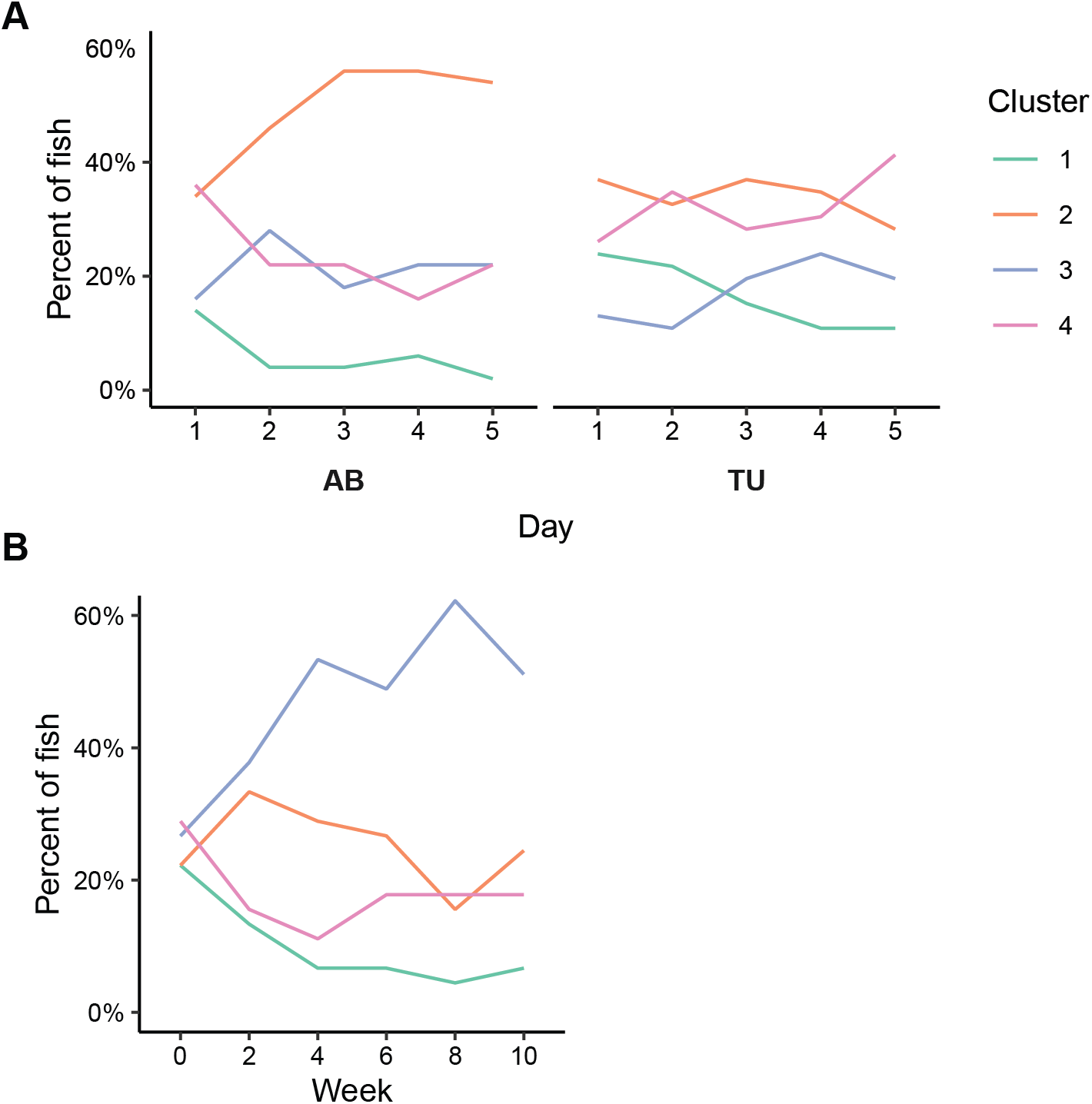
Percent of fish falling into each cluster during repeated exposures to the novel tank. **A**) Membership across five consecutive days for AB (left) and TU (right) fish. **B**) Cluster membership across biweekly (every other week) exposures of TU fish.

## Notes

### Competing Interest Statement

The authors have declared no competing interest.

### Summary of Updates

Added data testing consistency of behavioral clusters over 10 weeks (Figure 5) and updated methods/results/discussion accordingly.

